# Accurate Hippocampal Electric Field Modeling for Electroconvulsive Therapy: Numerical Accuracy and Convergence

**DOI:** 10.64898/2026.10.01.751407

**Authors:** Gregory M Noetscher, Derek Drumm, Esra Neufeld, Antonino Cassarà, Bryn Lloyd, Christopher C Abbott, Sergey N Makaroff, Zhi-De Deng

## Abstract

Accurate hippocampal electric field (E-field) prediction is important for individualized electroconvulsive therapy (ECT) dosing and emerging deep-brain stimulation approaches such as temporal interference. Because hippocampal fields are far weaker than those beneath the stimulation electrodes, they are highly sensitive to tissue interfaces, conductivity contrasts, and numerical discretization, so numerical error can rival the physical uncertainties of anatomy and conductivity. We investigate this numerical component of hippocampal dosimetry using individualized head models from eleven ECT patients. Identical *head40* segmentations, comprising 40 tissue compartments including the bilateral hippocampi, were simulated with three independent numerical methods: a charge-based boundary-element fast multipole method (BEM-FMM), a voxel-based hexahedral finite-element solver, and a conformal tetrahedral finite-element solver. For each, resolution was increased systematically and convergence assessed from the average hippocampal E-field magnitude. At the highest attainable resolutions, the three methods agreed closely, differing by 6%–7% in the left hippocampus and 9%–12% in the right. At default or coarse resolutions, however, they disagreed by 28.8% on average and up to 50.3% in the right hippocampus. Numerical convergence should therefore be established before differences are attributed to anatomical or conductivity uncertainty, and independent-solver agreement provides a practical basis for reliable hippocampal dosimetry.

## I. Introduction

Electroconvulsive therapy (ECT) remains the most effective acute treatment for severe, treatment-resistant, and psychotic depression, with remission rates substantially exceeding those of pharmacotherapy. Its clinical use is nonetheless limited by cognitive side effects, most prominently retrograde amnesia and reductions in verbal fluency, that prolong functional recovery and contribute to the treatment’s persistent stigma (Sackeim et al. 1993 [1], Sackeim et al. 2007 [2]). A large part of ECT technique refinement has been an effort to preserve antidepressant efficacy while reducing this cognitive burden, primarily through the choice of electrode placement and stimulus parameters. High-dose right unilateral (RUL) ECT, for example, can match the efficacy of bitemporal (BT) placement while producing less severe and less persistent memory impairment (Sackeim et al. 2008 [3]), and more focal configurations such as bifrontal (BF) placement have been introduced with similar goals (Kellner et al. 2010 [4]). Despite this, the clinician’s ability to predict the most effective and best-tolerated combination of placement, pulse width, and charge for an individual patient remains limited.

A central obstacle is that the conventional notion of ECT “dose”, the fixed extracranial current, typically set at 800 or 900 mA by the commercial device rather than by any biophysical rationale, is a poor proxy for the stimulation actually delivered to the brain. What drives both therapeutic and cognitive effects is the induced intracranial electric field (E-field), not the applied scalp current, and for a fixed current amplitude this field varies substantially between patients because intracranial dose depends on individual anatomy of skin, skull, cerebrospinal fluid, and brain tissue (Deng et al. 2014 [5]). This anatomical variability is especially pronounced in older adults, the population most often treated with ECT, and can simultaneously compromise efficacy (insufficient stimulation of mood-related circuitry) and safety (excessive stimulation of memory-related structures) (Deng et al. 2022 [6], Abbott et al. 2024 [7]). These observations have motivated a shift, led substantially by Abbott and colleagues, from fixed-amplitude dosing toward individualized, E-field–informed dosing, including amplitude-titration strategies in which pulse amplitude is set to achieve a target intracranial field rather than left fixed (Abbott et al. 2021 [8], Abbott et al. 2024 [7]). Realizing this approach depends directly on the ability to estimate, accurately and subject-specifically, the E-field delivered to the relevant deep-brain targets.

One such deep-brain target of interest is the hippocampus, which has been implicated in both ECT’s antidepressant mechanism and its cognitive side effects. A consistent body of imaging work has established that ECT produces measurable hippocampal change: Abbott et al. 2014 [9] linked hippocampal structural and functional changes to ECT response, and Takamiya et al. 2021 [10] localized the ECT-induced gray matter volume expansion predominantly to the medial temporal lobe, most pronounced on the right. The biophysical basis of this volume increase appears to be neurogenesis-independent; in a controlled animal model, Abe et al. 2024 [11] showed that the MRI-detectable hippocampal volume increase persisted when neurogenesis was inhibited and was instead accompanied by dendritic growth and increased synaptic density. Argyelan et al. 2019 [12] established the underlying dose–response, showing across 151 ECT-treated patients that regional E-field strength was associated with regional volume change in a dose-dependent fashion, with a particularly strong relationship in the hippocampus and amygdala and with individual regions expanding once the local field exceeded a threshold. Argyelan et al. 2021 [13] subsequently linked this hippocampal enlargement specifically to cognitive side effects, showing that the association between volume increase and cognitive impairment was hippocampal-specific, since volume changes in other regions correlated weakly or not at all with cognitive impairment. Deng et al. 2022 [6] connected these outcomes directly to the E-field, showing that right hippocampal E-field strength was linearly related to hippocampal neuroplasticity, which in turn mediated antidepressant response, while the field was also directly linked to a cognitive side effect (phonemic fluency decline), with roughly 112.5 V/m identified as the ceiling for cognitive safety. Loef et al. 2025 [14] recently replicated a similar relationship between regional E-field strength and cognitive side effects in an independent cohort, reinforcing that this variability is anatomically driven and clinically consequential. Together these findings imply a therapeutic window that must be hit within a given individual, which in turn requires knowing the actual field delivered to that individual’s hippocampus, precisely the quantity that accurate modeling must provide.

Accurate estimation of the hippocampal E-field is equally important for temporal interference (TI) stimulation (Grossman et al. 2017 [15]), an emerging noninvasive technique for steering a focal low-frequency envelope to deep brain structures. When the hippocampus is the intended target, relatively small anatomical differences can substantially alter both the amplitude and spatial distribution of the interference envelope. Errors in E-field modeling therefore translate into errors in the predicted location and magnitude of the modulation focus, potentially shifting stimulation onto adjacent structures such as the amygdala or parahippocampal gyrus rather than the targeted hippocampus. Computational modeling consequently plays a central role in optimizing electrode montages and verifying that the interference maximum is localized within the hippocampus while minimizing exposure of the overlying cortex. Recent work by Violante et al. 2023 [16] and Meng et al. 2025 [17] showed that hippocampal TI fields exhibit substantial inter-subject variability due to anatomical differences and correlations between those inter-subject dosimetric differences have been found to explain up to R^2^=40% of behavioral response variability [16], emphasizing that individualized E-field modeling is critical for reproducible targeting, treatment optimization, and meaningful interpretation of clinical outcomes. In contrast to ‘shallow’ cortical targets, whose sensitivity to modeling uncertainties has been examined in detail previously (Saturnino et al. 2019 [18], Cassarà et al. 2023 [19]), the hippocampus is a deep brain target at which the E-field strength may be 30-60 times lower than that immediately beneath the stimulation electrodes. Such relatively weak fields are highly sensitive to modeling accuracy, which deteriorates as current flows through intervening tissue compartments, especially if the relevant anatomy is highly heterogeneous and includes thin structures and tissue interfaces with high conductivity contrast.

This sensitivity raises a practical question: How can we trust modeling results obtained using a single finite-element solver with default or manually tuned parameters? It is part of simulation verification to ensure solver and discretization convergence [20] [21], but in some instances, solutions can converge to wrong results, as acknowledged by the IEEE IEC/IEEE 62704-1 standard for the validation of finite-differences time-domain (FDTD) simulators [22], which demands reproduction of expected grid-orientation-dependent numerical dispersion errors. If the underlying numerical solution is not sufficiently accurate, subsequent analyses of conductivity and/or segmentation uncertainties may be misleading, as the numerical error may be comparable to, or even exceed, the impact of more regularly quantified physical uncertainties (e.g., tissue properties, anatomical geometry, segmentation). The consequences are directly clinical: because ECT dosing is moving toward individualized, E-field–informed protocols, an unrecognized numerical error of tens of percent in the hippocampal field could push a patient outside the therapeutic window, either underdosing the antidepressant effect or exceeding the ceiling for cognitive safety. This study evaluates the numerical aspect of hippocampal field prediction accuracy using a dataset for eleven ECT patients. For this purpose, we will use three different numerical methods for solving the ohmic-current dominated electro-quasistatic equation (∇σ∇φ, where σ is the electrical conductivity, φ the electrical potential, and the electric field is obtained as E=−∇φ). The first is the boundary-element fast multipole method (BEM-FMM) operating directly with the underlying surface segmentation (Makarov et al. 2018 [23], Makarov et al. 2021 [24], Drumm et al. 2026 [25]). The second is the Sim4Life voxel-based finite-element method operating on a structured non-uniform hexahedral grid (Carbonell et al. 2026 [26]). The third is the Sim4Life finite element method operating with volumetric (conformal) tetrahedral mesh ([27]). For every method, we gradually increase the global (surface or volume) mesh resolution to investigate convergence and ensure that mutually consistent results are obtained.

## II. Materials and Methods

### 1. Subject models and ECT electrode placement

The anatomical models used in this study were derived from the patient cohort described in Deng et al. 2022 [6], to which the reader is referred for full details of recruitment, eligibility, ethical approval, and informed consent. Briefly, those data originate from a double-blind randomized clinical trial in patients with major depressive disorder, aged 50–80 years, who met clinical indication for ECT and were randomized to right unilateral (RUL) electrode placement at one of three pulse amplitudes (600, 700, or 800 mA). All procedures in the parent trial were conducted in accordance with relevant guidelines and regulations under the approval of the University of New Mexico Human Research Protections Office and all subjects provided written informed consent to the research protocol and study participation. From this cohort we selected T1/T2 1 mm isotropic MRIs and the corresponding electrode placements for 11 subjects. For each subject, 25 mm diameter electrodes were modeled in the RUL configuration, with other clinical placements (bifrontal, bitemporal) equally amenable to the same modeling approach. To isolate the numerical question addressed here, the montage and injected current were held fixed across all subjects: the current-controlled voltage electrodes were set at 800 mA. Fig. 1 shows a representative clinical ECT RUL model for one subject.

**Fig. 1:**
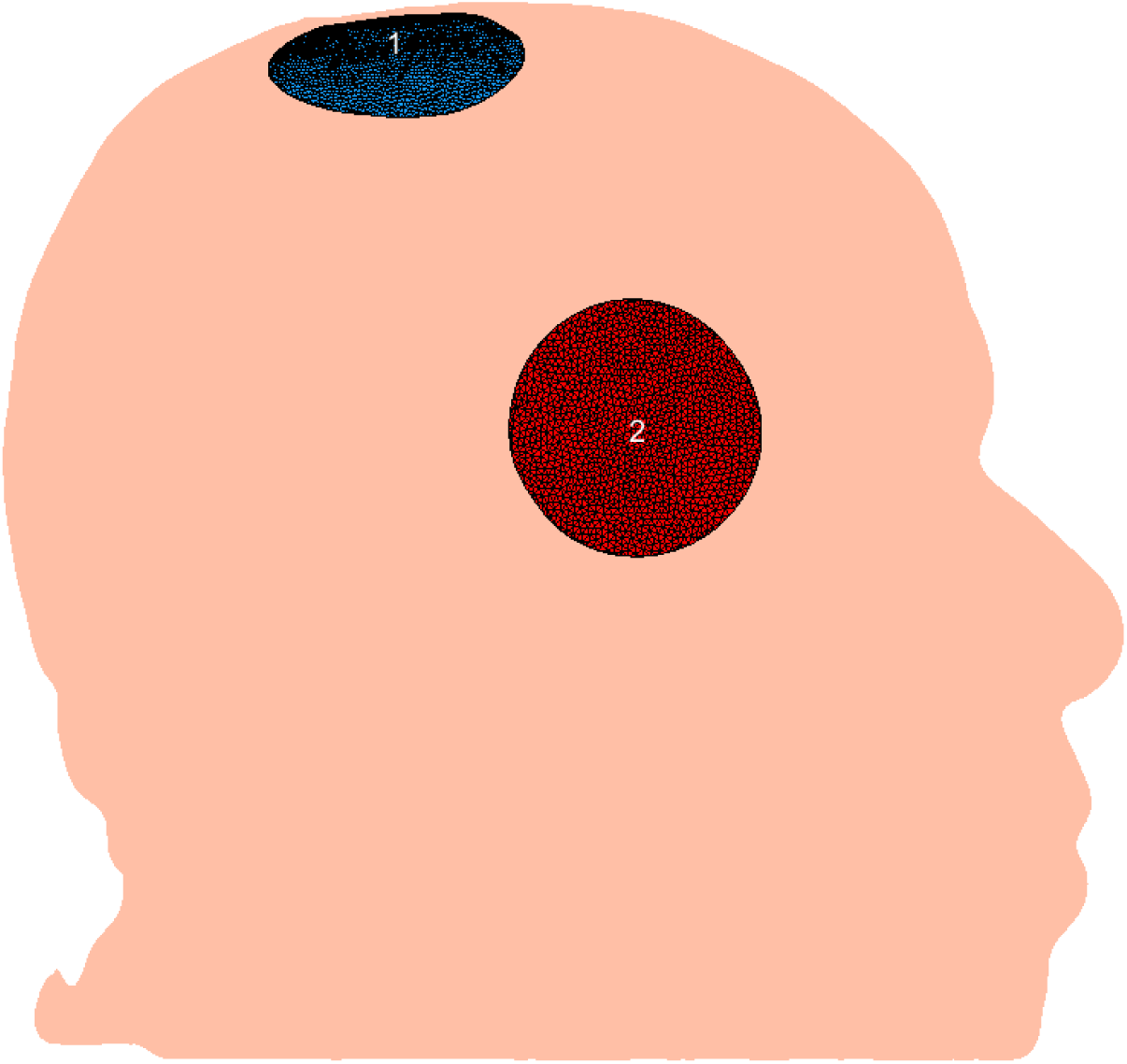
Sample ECT model with right unilateral electrode configuration

### 2. Segmentation pipeline

Surface meshes were extracted using the Sim4Life AI-based anatomy modeling pipeline *head40* ([28]) and assigned material properties from the low-frequency conductivity section of the IT’IS Tissue Properties Database (Baumgartner et al. 2025 [29]). *head40* identifies 40 distinct head tissue compartments including left and right hippocampi and surrounding deep brain structures as shown in Fig. 2. Sim4Life’s algorithm for surface extraction from segmentation label fields ensures that every interface between adjacent tissue compartments, except those at the domain boundary, is represented by a pair of exactly coincident but inversely oriented surface faces, with one face belonging to each adjacent compartment.

**Fig. 2:**
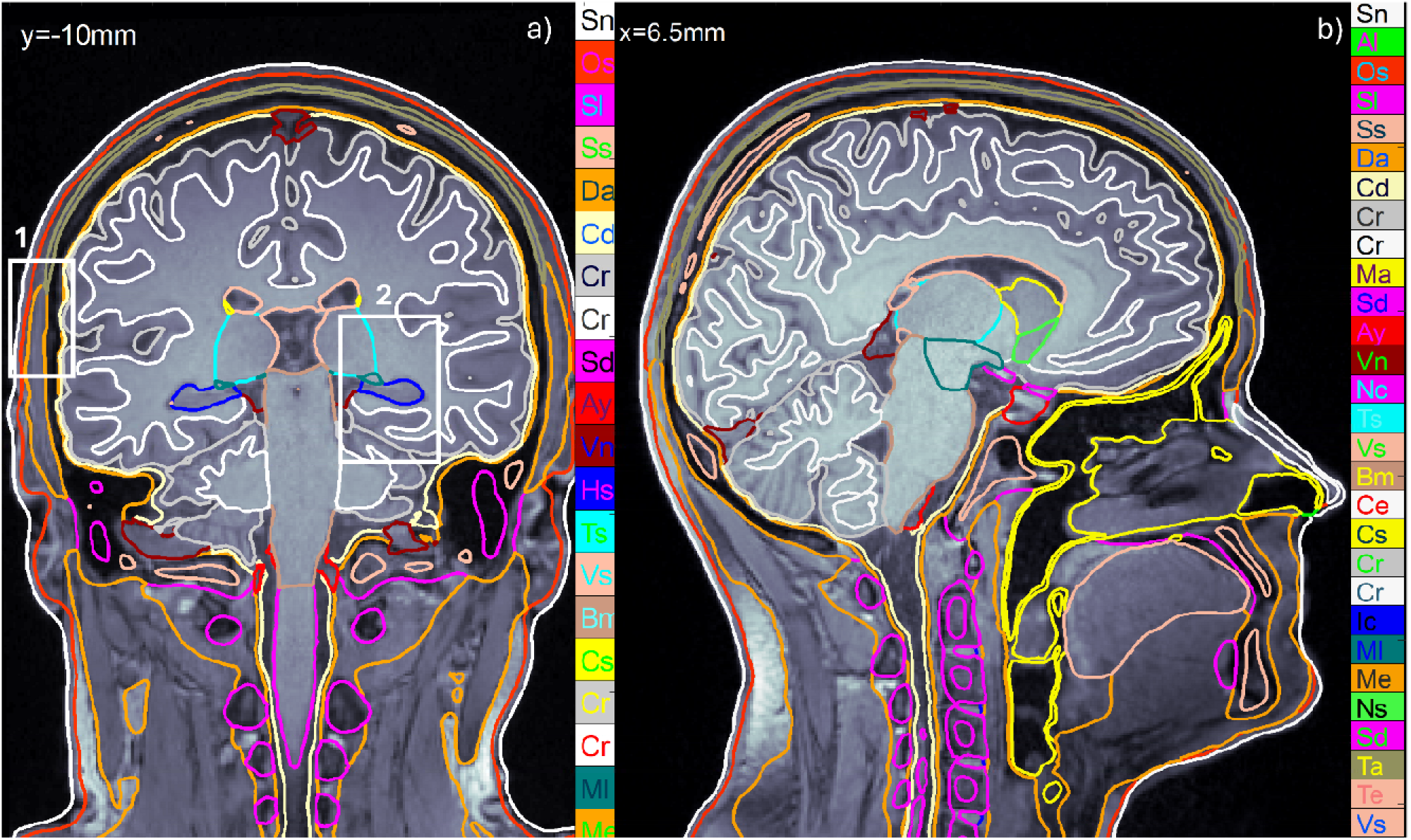
Example segmentation using the *head40* AI-assisted methodology from Sim4Life. Hippocampus cross-section is marked dark blue in the coronal plane.

### 3. Self-consistent non-onion segmentation

The resulting surface mesh is neither an “onion” model nor a manifold mesh (see Fig. 3, which shows two insets from Fig. 2 in detail). Instead, it provides a substantially more faithful representation of cranial anatomy than conventional nested (“onion”) models, such as SPM12/CAT/headreco (Saturnino et al. 2019 [30]) or the standard FreeSurfer (Fischl 2012 [31]), while remaining internally self-consistent, with no intersecting interfaces (Fig. 3). This property enables not only FEM but also BEM (boundary element method) analyses to be formulated unambiguously and solved with high numerical accuracy.

**Fig. 3:**
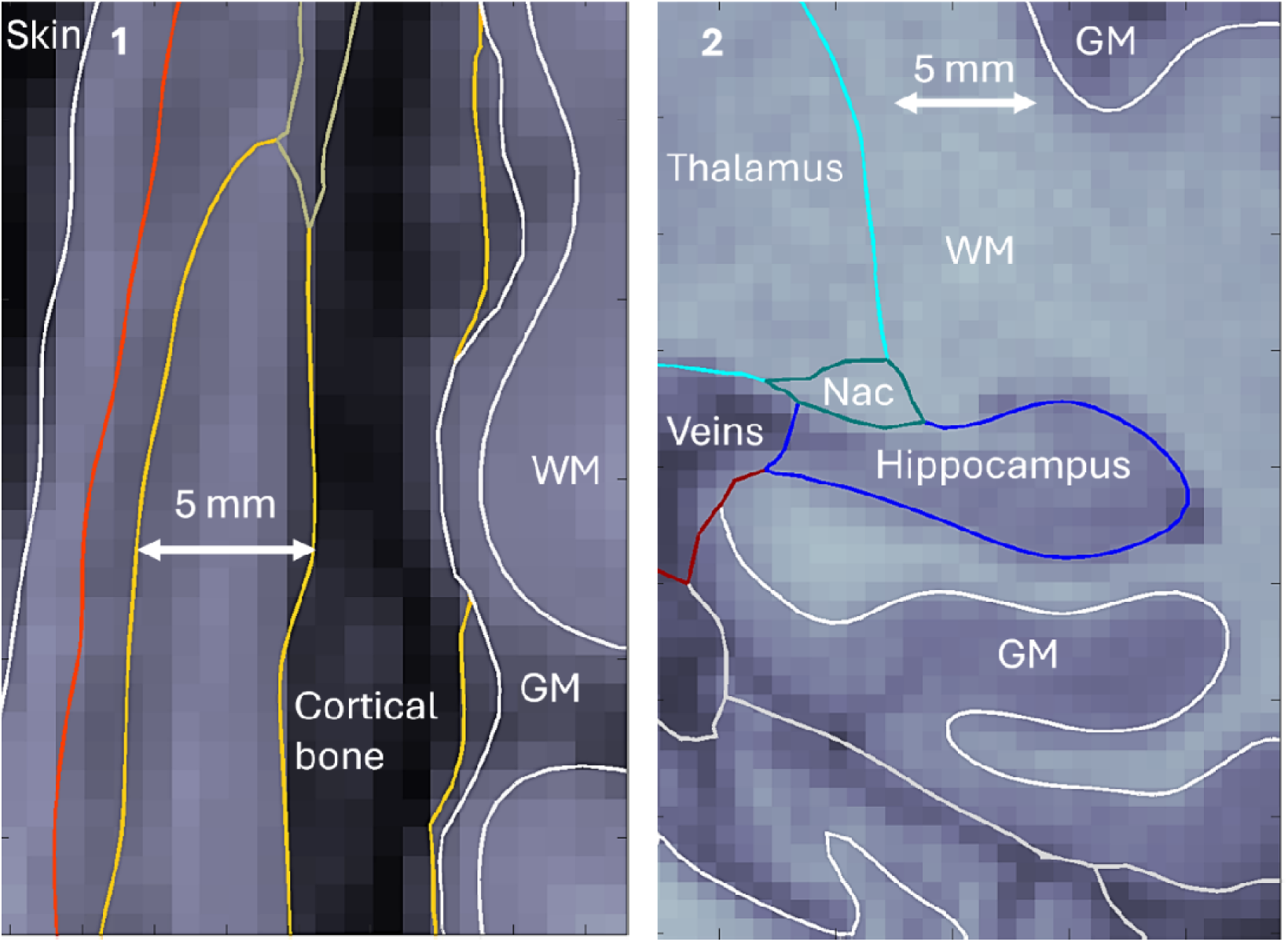
Detailed topology of *head40* segmentation at compartmental interfaces

For BEM applications, the duplicated interface facets are removed through a straightforward preprocessing step. The resulting *head40* model contains approximately 1 million facets in total, with an average edge length of 1.6–1.7 mm. The conductivity contrast associated with each remaining facet is then assigned according to the orientation of its outward normal vector, with corresponding inner/outer conductivities defined consistently with that orientation.

### 4. Observation/test points

For each subject, between 110,000–140,000 observation points were placed on a uniform grid strictly within the hippocampal volume (Fig. 4). We computed the average E-field magnitude for all observation nodes. Along with this, we also computed nodal vector fields and their 2-norm differences for the purposes of numerical evaluation.

**Fig. 4:**
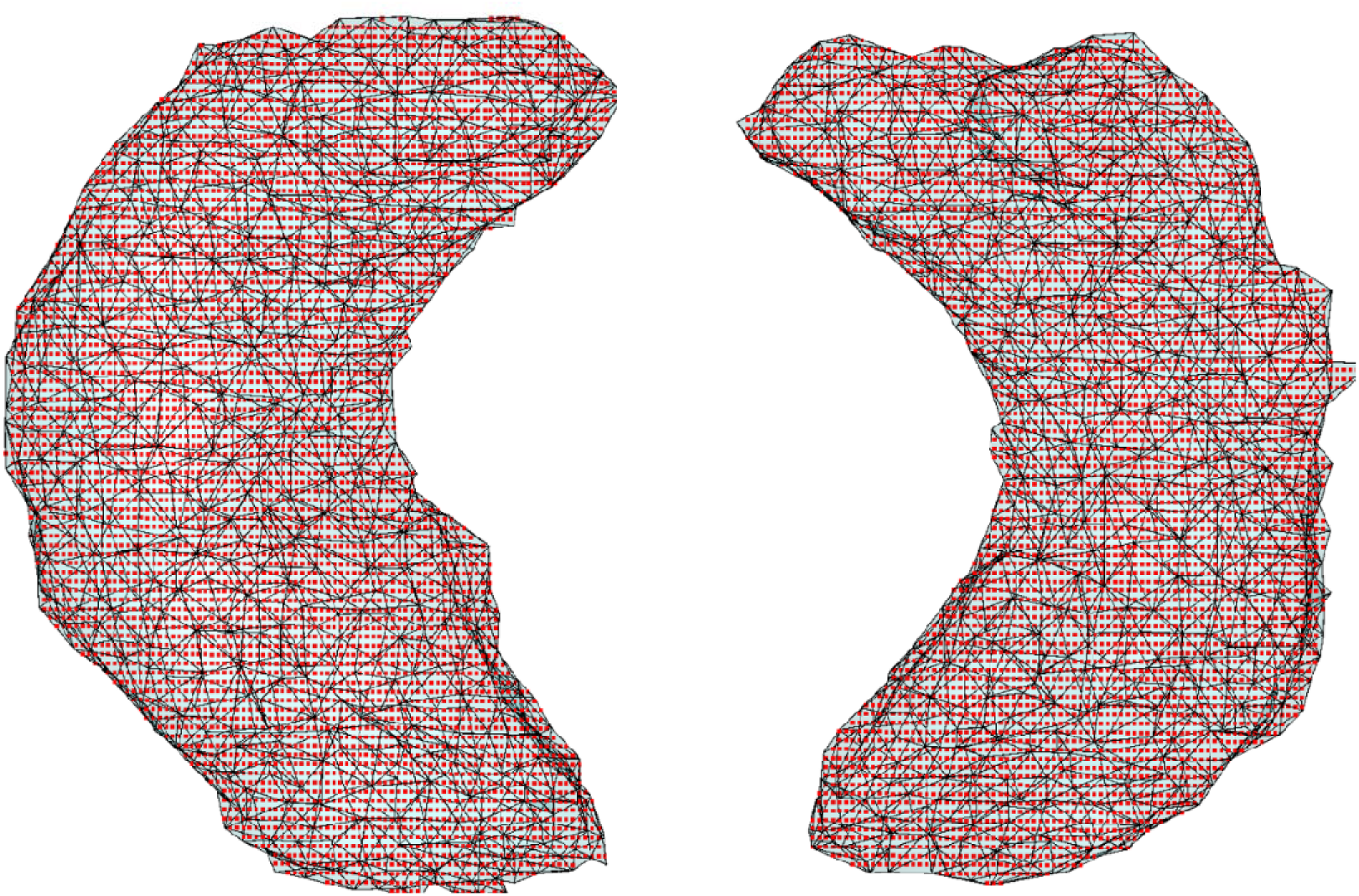
Typical observation points located within right and left hippocampal surface meshes. The number of points used was between 110k – 140k in total and varied by subject. Average electric field magnitude was found across all points for a given subject.

### 5. Numerical solvers

The charge-based BEM-FMM utilizes simplest zeroth-order (pulse) basis functions for the surface charge density at the interfaces and a sufficiently large nearfield size (Makarov et al. 2018 [23], Makarov et al. 2021 [24], Drumm et al. 2026 [25]), with approximately 1,000 neighboring facets for both the potential and the electric field. For these facets, the near-field integrals are computed using semi-analytical formulas instead of the standard FMM center-point approximation. The BEM solver also utilizes two surfaces meshes: the original *head40* model and the same model but subject to uniform 1:4 barycentric mesh refinement.

Given the known strong dependence of BEM-FMM solution accuracy on the number of neighbor integrals employed, a preliminary study of solution convergence with respect to the size of the nearfield was conducted. Fig. 5 demonstrates the importance of choosing the proper near-field size. There, typical convergence profiles for the average right (red) and left (blue) electric fields simulated with BEM-FMM in the hippocampus are presented for two patients from this study. For low values of the number of neighbor integrals employed, the solution is unreliable; as the number of integrals increases, the solution converges.

**Fig. 5:**
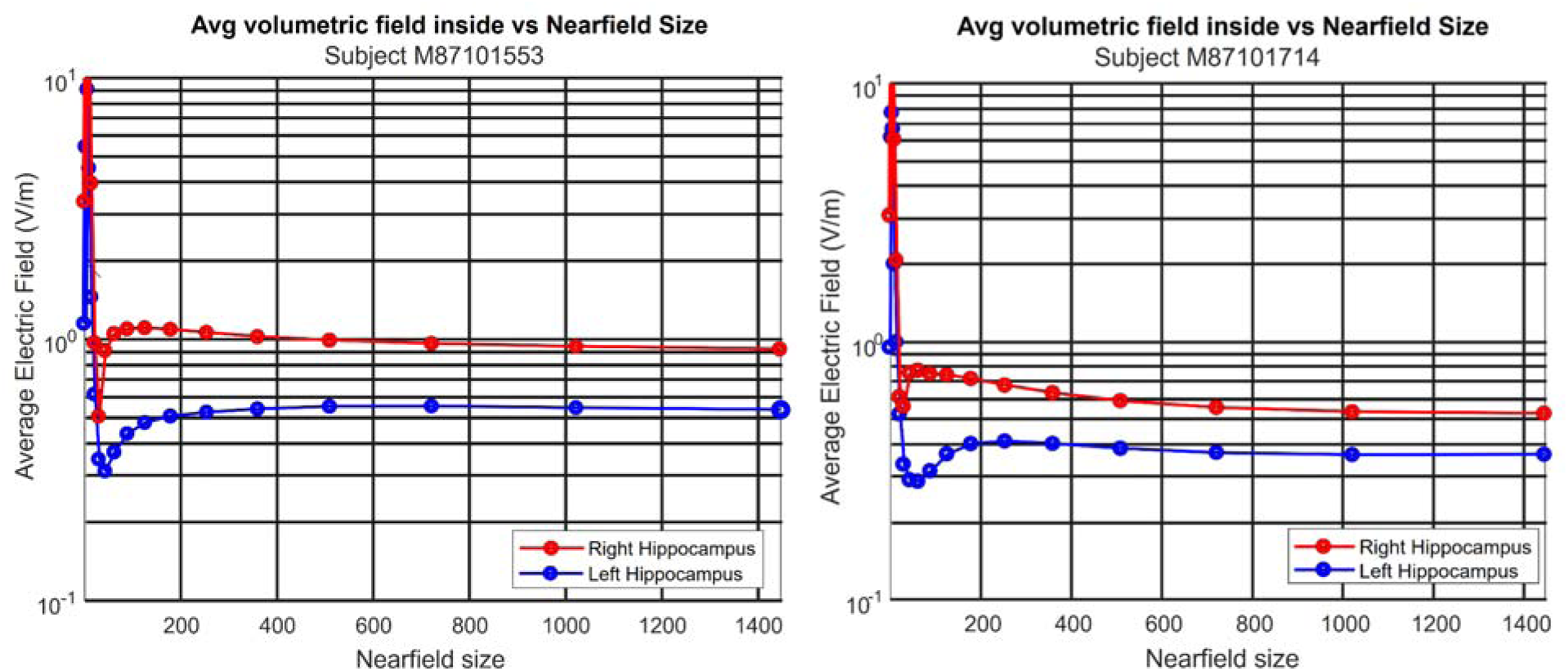
Typical convergence profile of the average right (red) and left (blue) electric fields simulated with BEM-FMM in the hippocampus for two patients employed in this study. A test problem with ± 1V voltage electrodes is considered. For low values of the number of neighbor integrals employed, the solution is unstable; as the number of integrals increases, the solution converges.

The voxel-based Sim4Life solver made use of successively refined uniform grids at spatial resolutions of 1 mm (default), 0.5 mm, and 0.25 mm. The Sim4Life FEM solver used successively refined tetrahedral meshing with maximum element edge lengths of 3 mm and 1.5 mm resolution. All three methods employed the identical *head40* base segmentation as well as the identical set of conductivity values (Baumgartner et al, 2025 [29]).

The grid resolution for each solution method was maximized to the fullest extent possible given the available computational hardware. This produced certain limitations on the size of the simulations and the number of elements employed. All Sim4Life computations (hexahedral and tetrahedral) were performed on a remote cluster running the Windows Server 2022 Datacenter operating system. This cluster employed Intel(R) Xeon(R) Gold 6542Y 2.90 GHz processors and had an available memory of 512 GB. For Sim4Life hexahedral simulations, the amount of memory on the cluster effectively limited the size of the simulation to approximately 1 billion hexahedra.. For Sim4Life tetrahedral simulations, the amount of memory on the cluster effectively limited the size of the simulation to approximately 200 M tetrahedra. A maximum edge length was enforced throughout the simulation space and the unstructured mesh around the electrodes was iteratively refined to fit within this limitation. All BEM-FMM computations were performed on a remote cluster running the Windows Server 2022 Datacenter operating system. This cluster employed Intel(R) Xeon(R) Platinum 8462Y+ 2.80 GHz processors and had an available memory of 1 TB. For BEM-FMM simulations with surface mesh sizes of approximately 1 M triangular elements, the number of neighbor triangles was effectively limited by cluster memory to 2048.

Finally, all simulations were performed with 1 V voltage-controlled electrodes. Because the problem is linear, the resulting fields can be scaled to any injected current *I*, and in the following we report all results at the clinically relevant ECT current of 800 mA.

## III. Results

### 1. Maps of cranial E-fields

Fig. 6 shows typical coronal/sagittal plane field maps for one subject obtained with the BEM-FMM method. There, E-field magnitude distribution (V/m) in the coronal (a) and sagittal (b) cross-sections passing through the center of mass of the right hippocampus (marked by a blue diamond) at 800 mA are shown. The colormap saturates at 820 V/m and a logarithmic field scale is used. The wide range of E-field strengths between electrodes (up to 1500 V/m in their immediate vicinity) and the left/right hippocampus (∼30–90 V/m) is highlighted for emphasis.

**Fig. 6:**
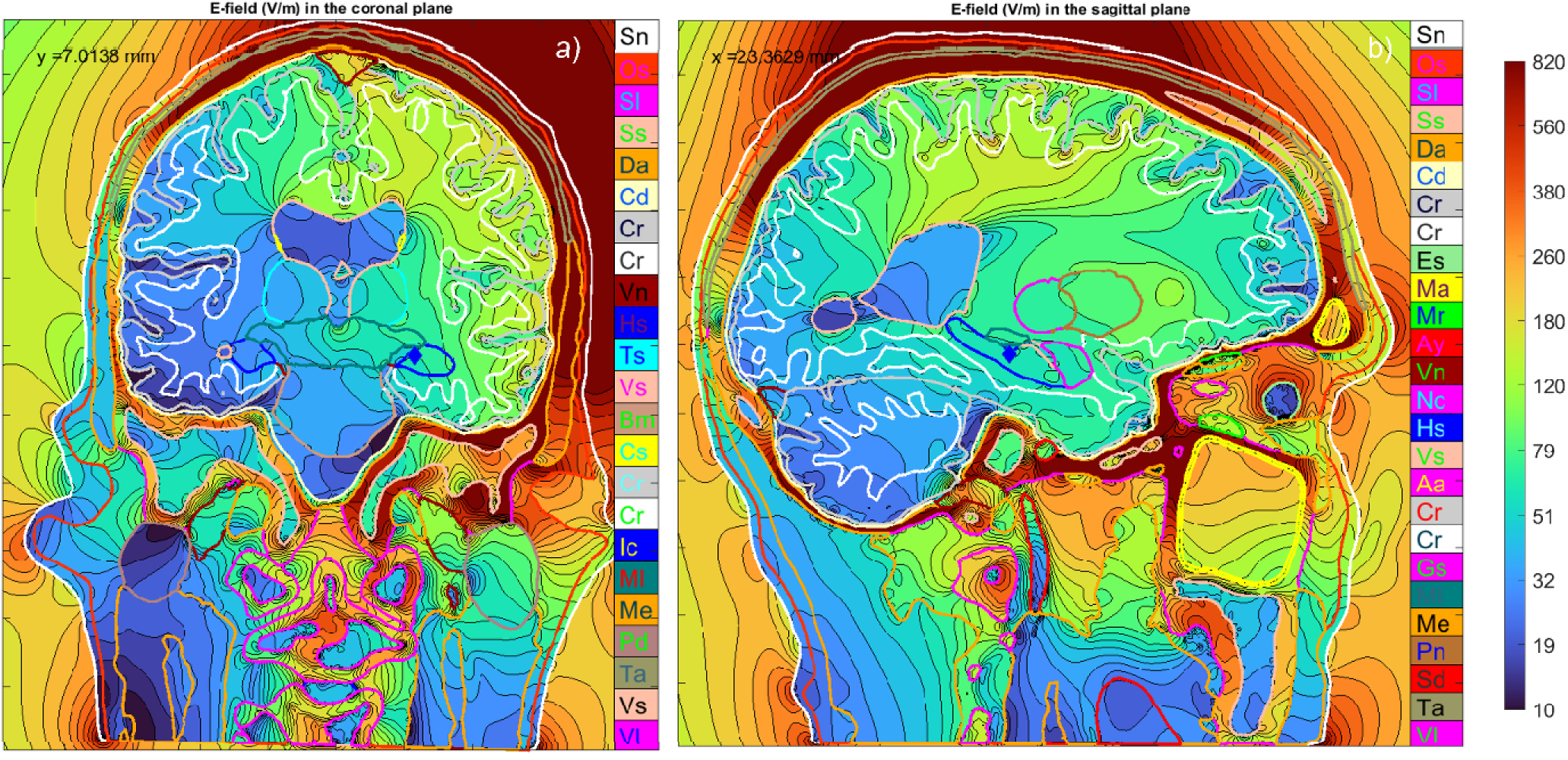
Simulated electric field in the coronal (a) and sagittal (b) planes passing through the center of mass of the right hippocampus, found using the BEM-FMM. The logarithmic color scale saturates at 820 V/m. Note the wide range of field magnitudes passing from the electrode (exceeding 1500 V/m) to the hippocampus (approximately 90 V/m).

### 2. Electrode currents for all three methods

Fig. 7 shows the net electrode currents for the ±1 V voltage-controlled voltage electrodes for all three methods and for all subjects. The BEM-FMM method uses the original *head40* mesh, the voxel-based Sim4Life solver uses 0.25 mm voxel size and the FEM Sim4Life solver uses 2 mm average mesh size. The average relative difference between the highest-resolution BEM-FMM and Sim4Life hexahedral solutions was 1.7%, and between BEM-FMM and the most refined Sim4Life tetrahedral solution, 1.2%. These differences directly reflect the difference in simulated impedance.

**Fig. 7:**
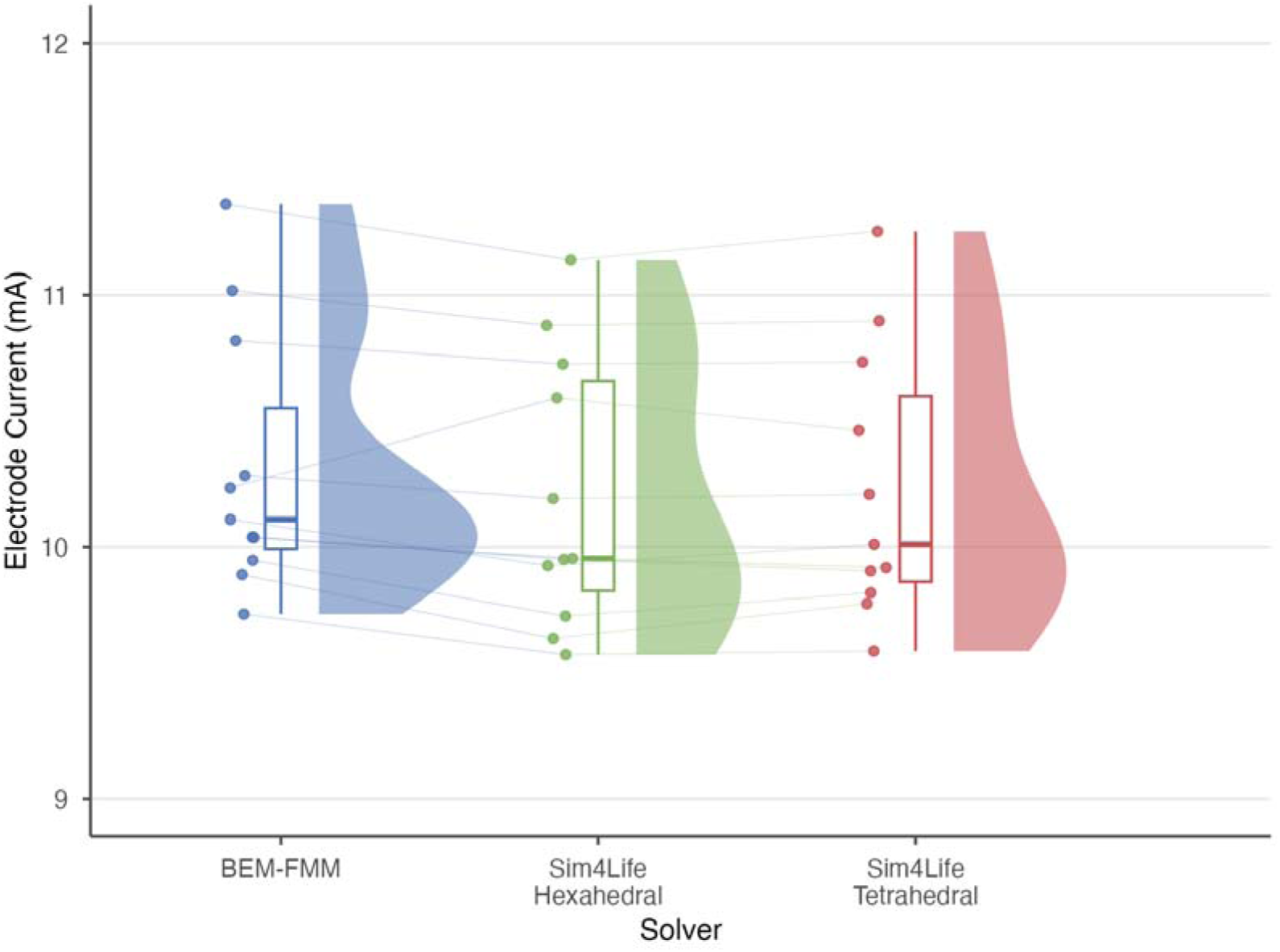
Net electrode currents computed for 1 V voltage-controlled electrodes across all subjects, using the BEM-FMM (blue), Sim4Life hexahedral (green), and Sim4Life tetrahedral (red) solvers. The relative difference between BEM-FMM and hexahedral results and the BEM-FMM and tetrahedral results were 1.7% and 1.2%, respectively.

### 3. Self-convergence results for the BEM-FMM solver

The whole-head model of approximately 1 million facets was processed using barycentric subdivision to produce a 1:4 mesh refinement and assess the impact of mesh size on BEM-FMM solution accuracy. With a greater number of triangles used to characterize mesh surfaces, fewer neighbor integrals are necessary to reach a converged field value: the original mesh used 1448 neighbor integrals, whereas the refined mesh used 512. As shown in Fig. 8, refinement increases the average hippocampal E-field. This effect is more pronounced in the right hippocampus (Fig. 8, right), most likely due to this structure’s proximity to the RUL electrode.

**Fig. 8:**
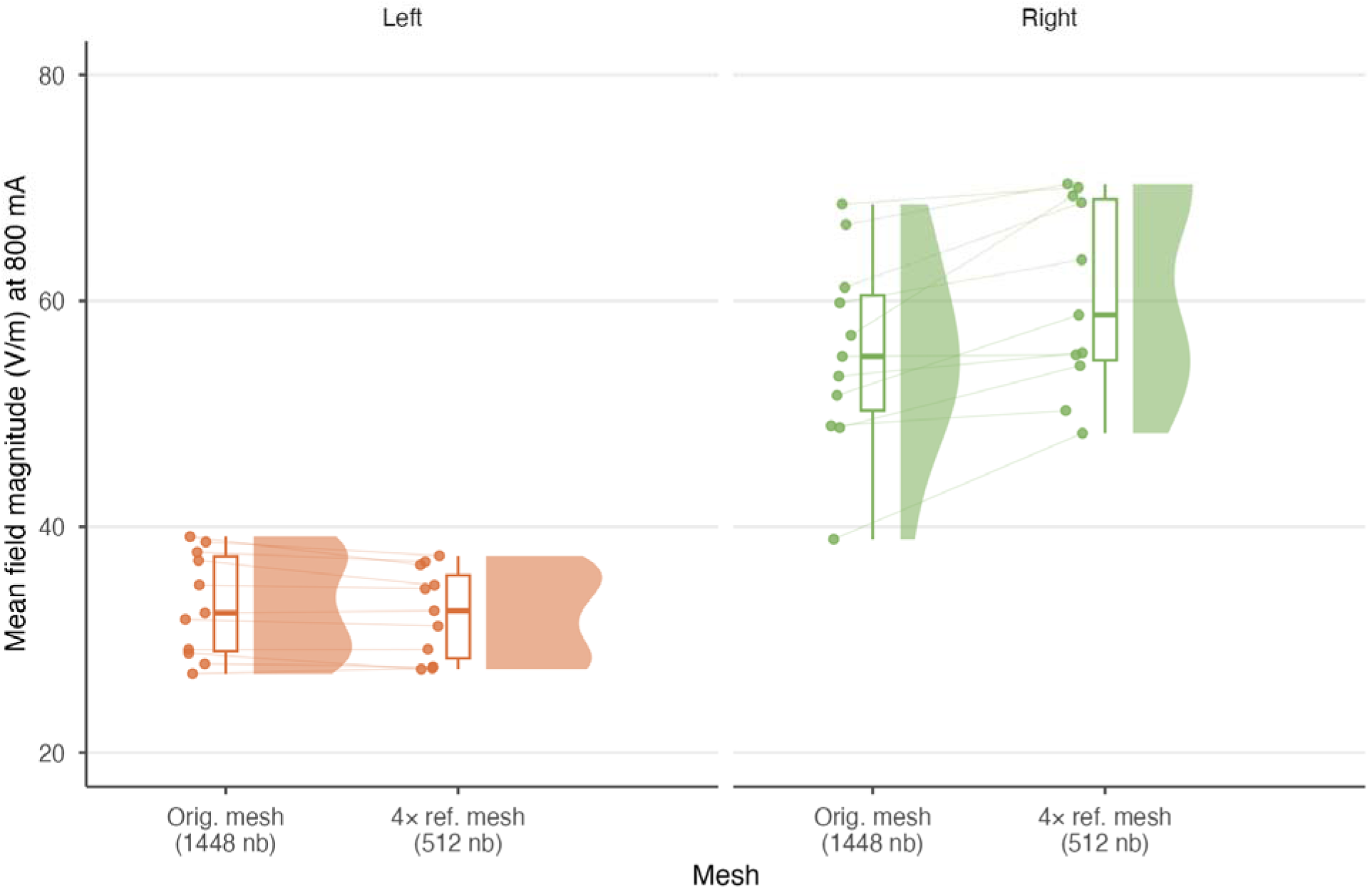
Effect of surface-mesh refinement on the mean hippocampal field magnitude (V/m at 800 mA) for the left and right hippocampus. Each panel compares the original *head40* surface mesh (1448 neighbor integrals) with the 4× barycentrically refined mesh (512 neighbor integrals); points are per-subject values, half-violins show the distribution, boxplots the summary statistics, and light lines connect the same subject across the two mesh conditions. Refinement increases the average field, an effect substantially larger in the right hippocampus, consistent with its proximity to the RUL electrode.

### 4. Self-convergence results for the Sim4Life voxel-based hexahedral solver

The accuracy of a solution computed with the Sim4Life hexahedral solver is dependent upon the number of hexahedral elements employed. As the number of elements increases, or the size of the voxel decreases, the simulation is expected to converge and provide the most accurate possible result. As shown in Fig. 9, the voxel edge length was decreased from 1 mm to 0.5 mm and again to 0.25 mm to generate three separate solutions, each with a uniform grid size over all regions of the simulation. Due to computational hardware limitations, 0.25 mm was maximum limit of this isometric scaling. Richardson extrapolation was used to estimate an infinitely resolved solution based on the finite resolution results. The values found via extrapolation continue the trend displayed by the lower resolution values, i.e., a finer resolution reduces the average value of the E-field in both the right and left hippocampus regions. This trend is more visible in the right hippocampus (Fig 9, right) than the left and may be due to higher field values.

**Fig. 9:**
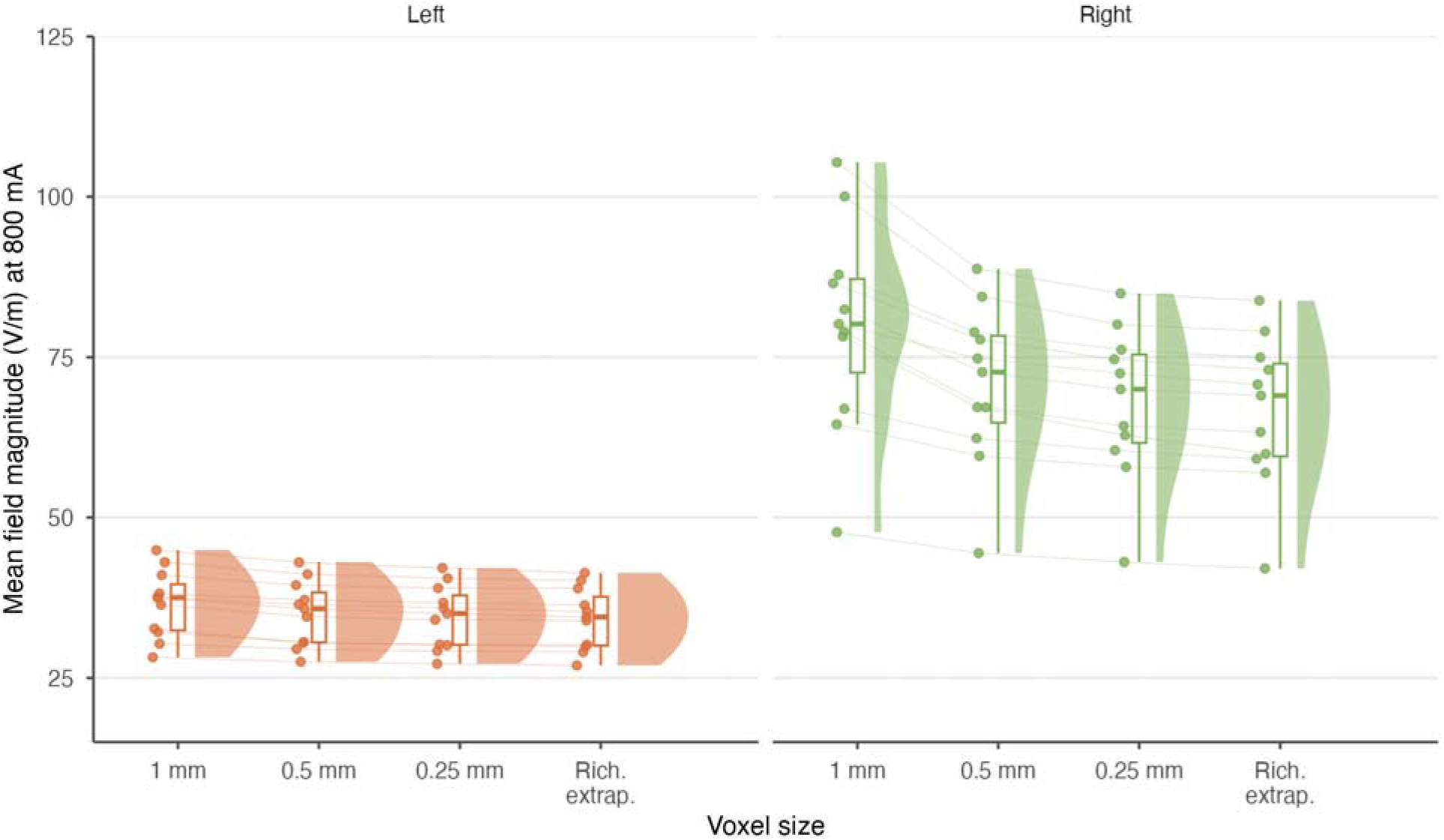
Convergence of the Sim4Life hexahedral solution for the mean hippocampal field magnitude (V/m at 800 mA), left and right hippocampus. Each panel shows the field at voxel edge lengths of 1 mm, 0.5 mm, and 0.25 mm, together with the Richardson extrapolation to the continuum limit; points are per-subject values, half-violins the distribution, boxplots the summary statistics, and light lines connect the same subject across resolutions. Finer resolution decreases the average field, with the extrapolation continuing this trend; the effect is substantially larger in the right hippocampus.

### 5. Self-convergence results for Sim4Life’s Unstructured FEM solver

The convergence of the Sim4Life tetrahedral solution was assessed by comparing two global mesh resolutions, with element edge lengths of 3 mm and 1.5 mm throughout the simulation region (Fig. 10). Consistent with the hexahedral results, refinement reduces the average hippocampal field, and the effect is more pronounced in the right hippocampus (Fig. 10, right) than the left. The change between the two resolutions is modest, however, indicating that the tetrahedral solution is already close to convergence at the coarser 3 mm resolution.

**Fig. 10:**
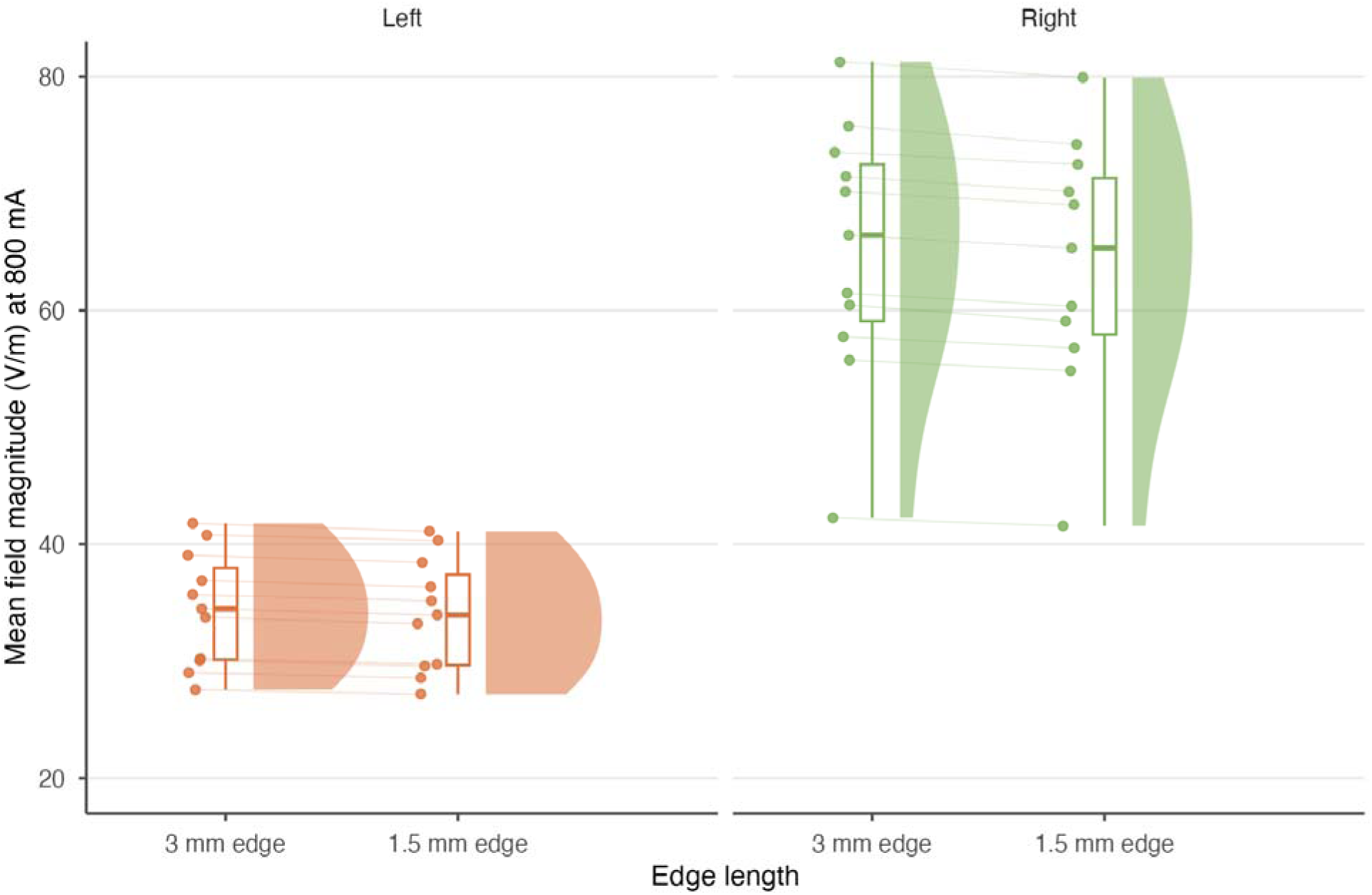
Convergence of the Sim4Life tetrahedral solution for the mean hippocampal field magnitude (V/m at 800 mA), left and right hippocampus. Each panel compares global element edge lengths of 3 mm and 1.5 mm; points are per-subject values, half-violins the distribution, boxplots the summary statistics, and light lines connect the same subject across the two resolutions. Finer resolution decreases the average field, an effect that is larger in the right hippocampus and modest overall.

### 6. Comparison of the three solvers at best resolution

Fig. 11 compares the three solvers, each at its finest attainable resolution: BEM-FMM (refined surface mesh, 512 neighbor integrals), the Sim4Life hexahedral solution obtained by Richardson extrapolation, and the Sim4Life tetrahedral solution at 1.5 mm. The three methods agree closely in the left hippocampus, whereas the right hippocampus shows greater inter-method variability. Across both structures the BEM-FMM tends to yield somewhat lower fields than either Sim4Life solver, most visibly in the right hippocampus.

**Fig 11:**
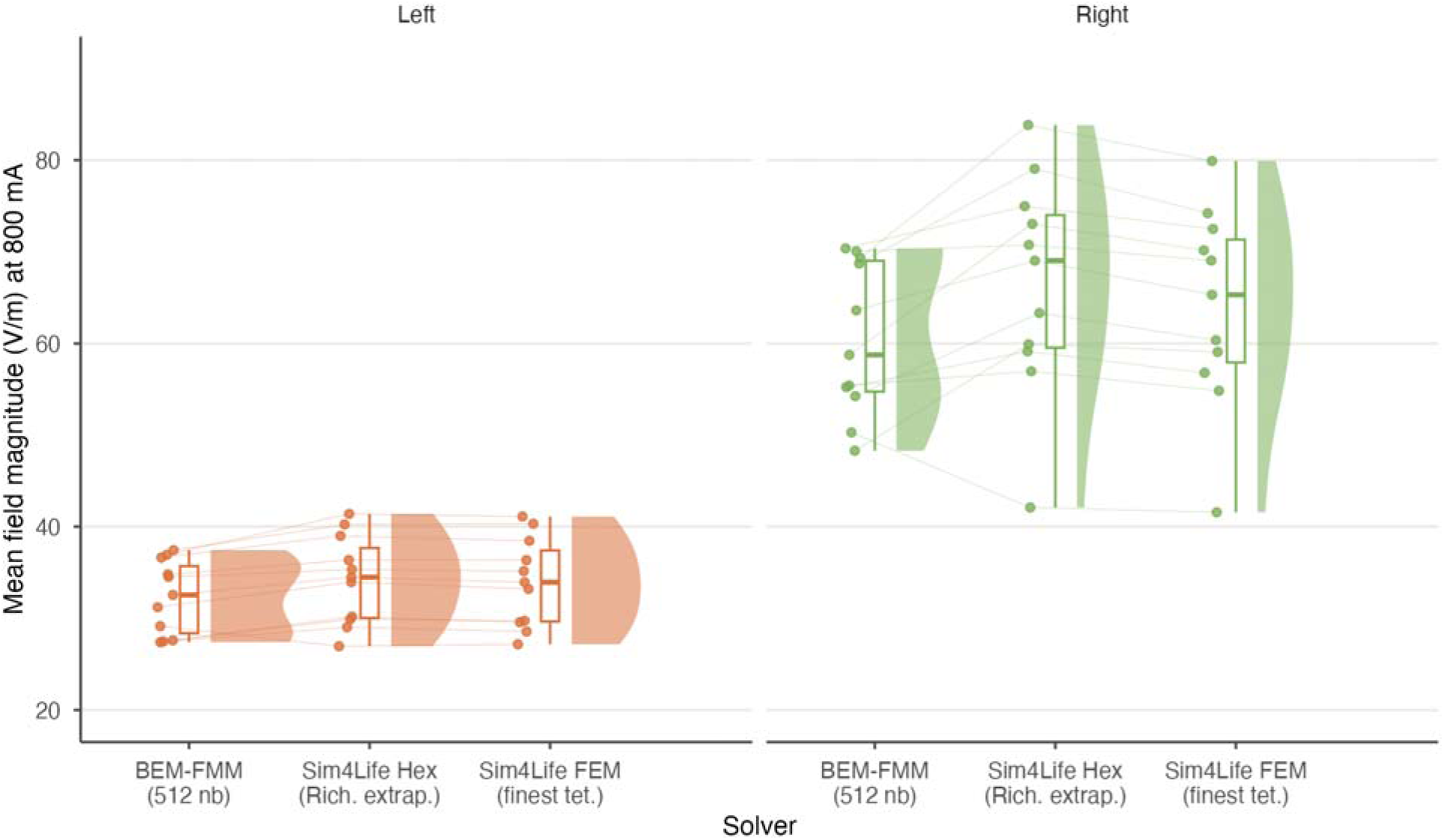
Comparison of the three solvers at their best attainable resolution for the mean hippocampal field magnitude (V/m at 800 mA), left and right hippocampus. Each panel shows BEM-FMM (refined mesh, 512 neighbor integrals), the Sim4Life hexahedral solution (Richardson extrapolation), and the Sim4Life tetrahedral solution (finest resolution); points are per-subject values, half-violins the distribution, boxplots the summary statistics, and light lines connect the same subject across solvers. Agreement is close in the left hippocampus and more variable in the right, where the BEM-FMM values tend to fall below both Sim4Life solutions.

Quantitatively, all relative differences are reported as the mean across the eleven subjects of the per-patient absolute relative difference between two solutions. Between the highest-resolution BEM-FMM and the finest Sim4Life tetrahedral solution, the relative differences in right and left hippocampal field magnitude were 8.8% and 5.8%, respectively, and between BEM-FMM and the Richardson-extrapolated Sim4Life hexahedral solution, 11.6% and 6.8%. These converged differences stand in sharp contrast to those obtained at default or coarse settings: the BEM-FMM with the default surface segmentation and the hexahedral Sim4Life solver at its default 1 mm isometric grid differ by 28.8% on average in the right hippocampus, reaching 50.3% in the worst individual case. Averaged over all subjects, the mean hippocampal field magnitudes at 800 mA in the right and left hippocampus were 60.4 and 32.3 V/m for the BEM-FMM, 66.6 and 34.3 V/m for the Richardson-extrapolated hexahedral solution, and 64.0 and 34.0 V/m for the finest tetrahedral solution.

## IV. Discussion

At the finest resolutions attainable with the available hardware, the three solvers produced closely consistent hippocampal fields, agreeing to within roughly 6%–7% in the left hippocampus and 9%–12% in the right (Fig. 11). The consistently smaller differences in the left hippocampus, and the tendency of the BEM-FMM to predict somewhat lower fields than either Sim4Life solver, are both visible directly in Fig. 11 and are consistent with the right hippocampus lying closer to the RUL electrode and carrying the higher, more resolution-sensitive fields. That three independent formulations, a charge-based boundary-element method and two finite-element discretizations on structurally different grids, converge to comparable values provides stronger evidence of accuracy than the self-convergence of any single solver, since the methods share neither their discretization of the geometry nor their sources of numerical error. Agreement between independent formulations is therefore a useful practical criterion for establishing confidence in deep-brain dosimetry.

These levels of agreement were only obtained because each method was pushed to the fullest extent the hardware allowed. Had the solvers instead been run at their default or coarse settings, the discrepancies would have been far larger, reaching 28.8% on average and 50.3% at worst in the right hippocampus. This gap between converged and default-resolution results is the central practical finding of the study, and it highlights the need for adaptive mesh refinement development, which concentrates mesh or grid resolution in the limited regions that most influence the deep-brain solution rather than incurring the memory and time cost of globally uniform refinement. Such approaches have recently been developed for the BEM-FMM (Drumm et al. 2026 [25], Wartman et al. 2024 [32], Lee et al. 2018 [33]), and further work in this direction is likely to be necessary.

The magnitude of this numerical error is best appreciated by comparison with the physical uncertainties that hippocampal dosimetry studies more commonly investigate. Conductivity uncertainty, for example, has been estimated to produce a standard deviation in field magnitude on the order of 20%–25% of the mean for surface-electrode montages (Saturnino et al. 2019 [6]). The converged inter-solver differences found here, of roughly 6%–12%, are smaller than this conductivity uncertainty, whereas the coarse-resolution numerical error of nearly 30% is comparable to or larger than it. In other words, a hippocampal field computed on a default mesh can carry a numerical error that rivals or exceeds the conductivity-driven uncertainty the study set out to characterize. It implies that numerical convergence must be established before differences in predicted hippocampal fields are attributed to anatomical, segmentation, or conductivity uncertainty; otherwise, the two are confounded.

The right hippocampus was consistently more sensitive to numerical resolution than the left, across all three solvers and both the mesh-refinement and voxel-convergence analyses. Three mutually reinforcing factors account for this. It lies closer to the right unilateral electrode, so it sits in a region of stronger and more rapidly varying field; it carries the higher absolute field magnitude, roughly twice that of the left, so a given fractional discretization error translates into a larger absolute deviation; and the current reaching it passes through nearby regions of high conductivity contrast, including the skull, where potential gradients are steep and discretization error accumulates. The same reasoning implies that the sensitivity of a deep target to numerical resolution is not fixed but depends on its position relative to the electrodes, and that targets near strong-gradient, high-contrast pathways warrant the most careful convergence testing.

An interesting feature evident across all subjects is a region of high field magnitude just below the right side of the cranial cavity, seen in Fig. 6, which is somewhat unexpected given the RUL electrode placement. This appears to be the result of large amounts of surface charge accumulation at the boundaries of the skull, which have lower conductivity values relative to the surrounding tissues; this is especially true of skull cortical bone, which was assigned a conductivity of 6.3 mS/m. Considering the size of these areas and the strength of the E-field, their influence on the right hippocampal field is likely to be appreciable. These areas of high potential gradients, especially across media of high conductivity contrast, therefore require special segmentation considerations where deep brain targets are concerned.

These numerical considerations bear directly on the individualized, E-field–informed dosing toward which ECT and deep-target stimulation are moving. If pulse amplitude is to be titrated to place the hippocampal field within a therapeutic window, for example below the cognitive-safety ceiling of roughly 112.5 V/m identified by Deng et al. 2022 [6], then a numerical error of tens of percent in the predicted field is not a technical detail but a clinically material one, potentially the difference between an effective and a cognitively harmful dose. Reliable subject-specific dosimetry therefore requires that the numerical solution be demonstrably converged before it is used to inform amplitude selection.

Several limitations should be noted. First, this study isolated the numerical component of hippocampal field prediction while holding the segmentation and conductivity values fixed; it therefore characterizes one part of the total dosimetric uncertainty rather than the whole, and the conductivity uncertainty referred to above is treated only for comparison, not quantified here. Second, convergence was limited by available hardware rather than carried to the true asymptotic limit: the hexahedral solver reached 0.25 mm, the tetrahedral solver 1.5 mm, and the BEM-FMM 512 neighbor integrals on the refined mesh, and the hexahedral continuum estimate relied on Richardson extrapolation. Third, all comparisons used a single fixed montage and injected current (RUL, 800 mA); other clinically used placements and amplitudes were not modeled here, although the same numerical framework applies to them directly. Finally, for the coarse-resolution comparison the field at each refinement level was scaled to 800 mA using the solver’s converged electrode current rather than the current drawn by the coarse model itself, so the absolute coarse-mesh field values are approximate; this does not affect the convergence trends or the converged-resolution differences, which use each solution’s own current.

## V. Conclusions

Accurate prediction of hippocampal E-fields requires careful attention to numerical discretization alongside the anatomical and conductivity uncertainties more commonly examined. Applying three independent numerical formulations, a charge-based BEM-FMM, a voxel-based hexahedral FEM, and a conformal tetrahedral FEM, to identical high-resolution *head40* models of eleven ECT patients, we found that all three converge to comparable hippocampal fields when sufficiently refined, agreeing to within roughly 6%–7% in the left hippocampus and 9%–12% in the right. At default or coarse resolutions, by contrast, the same models disagreed by 28.8% on average and by as much as 50.3% in the right hippocampus. Because these methods share neither their discretization of the geometry nor their sources of numerical error, their agreement at converged resolution provides a stronger basis for confidence in deep-brain dosimetry than the self-convergence of any single solver. Errors of the size seen at coarse resolution, however, are comparable to or larger than the physical uncertainties that hippocampal dosimetry studies typically set out to characterize, so numerical convergence must be established before differences in predicted fields are attributed to anatomy, segmentation, or conductivity. Because global uniform refinement to convergence is computationally expensive, and because deep targets near the electrodes and along steep conductivity contrasts, such as the right hippocampus here, are the most demanding, these findings also motivate development in adaptive mesh refinement that concentrates resolution where it most affects the deep-brain solution.

For ECT and other deep-brain stimulation modalities now moving toward individualized, E-field-informed dosing, such numerical validation is an essential prerequisite. Only once the computed field is demonstrably converged can it be used to place the hippocampal dose within its therapeutic window, and thus to base stimulation on the field actually delivered to the target rather than on the applied scalp current alone.

## Acknowledgements

This study has received support from the NIH grants R01MH130490 and R01EB035484 (SNM) as well as the Intramural Research Program of NIH, NIMH ZIAMH002955 (Z-DD).

## Author Contributions and Declaration of Interest

G.M.N. and S.N.M. conceived the study and designed the overall research framework. G.M.N., S.N.M., and Z.-D.D. developed and implemented the computational modeling approach. G.M.N., A.C., E.N., and B.L. contributed to data processing, numerical analysis, and validation of Sim4Life simulations. G.M.N. and S.N.M contributed to data processing, numerical analysis, and validation of BEM-FMM simulations. D.D. provided analysis and validation of BEM-FMM and BEM-FMM augmented with adaptive mesh refinement. Z.-D.D. and C.C.A. contributed to interpretation of the results in the context of brain stimulation, neuroimaging, and neurophysiology. G.M.N and S.N.M. led data interpretation and wrote the manuscript. All authors contributed to discussion of the results and provided critical feedback on the manuscript.

## Data Availability

The hippocampal fields obtained with the three numerical methods are made available in *.csv format

